# Soluble guanylyl cyclase, the NO-receptor, regulates endothelium-dependent vascular relaxation via its transnitrosation activity

**DOI:** 10.1101/2024.10.28.620487

**Authors:** Waqas Younis, Chuanlong Cui, Tanaz Sadeghian, Pia Burboa, Ping Shu, Yong Qin, Lai-Hua Xie, Mauricio Lillo Gallardo, Annie Beuve

**Affiliations:** Department of Pharmacology, Physiology and Neuroscience, New Jersey Medical School at Rutgers, Newark, NJ 07103; Department of Surgery, New Jersey Medical School at Rutgers, Newark, NJ 07103; Department of Cell Biology & Molecular Medicine, New Jersey Medical School at Rutgers, Newark, NJ 07103

**Keywords:** Soluble guanylyl cyclase, Connexin 43, S-nitrosation, mesenteric arteries, vascular resistance

## Abstract

We previously demonstrated that the NO-receptor soluble guanylyl cyclase (GC1) has the ability to transnitrosate other proteins in a reaction that involves, in some cases, oxidized Thioredoxin 1 (oTrx1). This transnitrosation cascade was established *in vitro* and we identified by mass spectrometry and mutational analysis Cys 610 (C610) of GC1 α-subunit as a major donor of S-nitrosothiols (SNO). To assay the relevance of GC1 transnitrosation under physiological conditions and in oxidative pathologies, we studied a knock-in mouse in which C610 was replaced with a serine (KI αC^610S^) under basal or angiotensin II (Ang II)–treated conditions. Despite similar GC1 expression and NO-stimulated cGMP-forming activity, the Ang II-treated KI mice displayed exacerbated oxidative pathologies including higher mean arterial pressure and more severe cardiac dysfunctions compared to the Ang II-treated WT. These phenotypes were associated with a drastic decrease in global S-nitrosation and in levels of SNO-Trx1 and SNO-RhoA in the KI mice. To investigate the mechanism underlying the dysregulation of blood pressure despite an intact NO-cGMP axis, pressure myography and *in vivo* intravital microscopy were conducted to analyze the vascular resistance tone. Both approaches indicated that, even in the absence of oxidative stress, the single mutation C610S led to a significant deficiency in acetylcholine-induced vasorelaxation while smooth muscle relaxation in response to NO remained unchanged. These findings indicate that the C610S mutation uncoupled the two NO signaling pathways involved in the endothelium and smooth muscle vasorelaxation and suggest that GC1-dependent S-nitrosation is a key player in endothelium-derived hyperpolarization.

## INTRODUCTION

Cardiovascular health is highly dependent on nitric oxide (NO) signaling and is extensively compromised by oxidative stress. Soluble guanylyl cyclase (GC1) is a heme-containing heterodimer, which catalyzes the production of cGMP upon stimulation of NO binding to the heme. The NO-GC1-cGMP pathway is critical to smooth muscle relaxation of blood vessels, as well as protecting the heart from myocardial infarction, fibrosis and negatively modulating cardiac contractility(1-3). In fact, GC1 is currently one of the most sought-after targets to treat cardiovascular diseases(4). NO also signals in the cardiovascular system by a non-canonical pathway, S-nitrosation (also called S-nitrosylation), which is the addition of a NO moiety to a cysteine (Cys)(5). As other post-translational modifications (PTMs), S-nitrosation changes protein function, localization or interaction.

We recently discovered that GC1 mediates as well the NO non-canonical pathway. We showed in cardiac and smooth muscle cell lines that GC1 has the ability to drive S-nitrosation of over 200 proteins via a transnitrosation activity, i.e., the transfer of S-nitrosothiol groups (SNO) through specific protein-protein interactions (6). GC1 transnitrosation activity is favored under oxidative stress, a condition that could lead to disruption of the NO-GC1-cGMP by oxidation of the heme and thiols of GC1, reduction of NO availability and increased S-nitrosation levels in cells(6-8). Interestingly, Thioredoxin 1 (Trx1) a major regulator of cellular thiol redox is one of the GC1 transnitrosation targets. The transnitrosation reaction was unidirectional from GC1 to Trx1 and took place with oxidized Trx1 (oTrx1), which acquires a transnitrosation activity when oxidized(9) (10, 11). We showed that SNO-GC1 “uses” oTrx1 as a SNO relay to amplify the number of SNO-targets. In fact, using Angiotensin II (AngII) as an oxidative/nitrosative inducer in cardiac cells, we discovered that SNO-GC1/oTrx1 complex S-nitrosates the small GTPase RhoA, in turn inhibiting its activity (6). Using Mass spectrometry, we identified Cys 610 (C610) of the α subunit of GC1 as the main SNO donor and Cys73 (C73) of Trx1 as the main SNO recipient/acceptor. We confirmed the crucial role of αC610 in GC1 transnitrosation activity by mutational analysis in smooth muscle cells. Of importance, the NO-stimulated cGMP-forming activity per se was not affected in the mutant. (12)

To determine the (patho)physiological significance of GC1 transnitrosation activity in the cardiovascular system, we created a knock-in mouse for which αC610 was replaced by a serine (KI αC^610S^). We used chronic infusion of AngII, which is a well-described model to increase blood pressure and to induce cardiac hypertrophy and electrical dysfunction. We previously showed that oxidative stress generated by Ang II infusion induces S-nitrosation in rodent tissues, and affects vascular reactivity and cardiac function (13, 14). Together with the idea that S-nitrosation could be cardioprotective (15, 16) and that it modulates several signaling pathways involved in vascular reactivity, including calcium signaling (17, 18), we sought to determine whether disruption of the transnitrosation cascade in the knock-in (KI) mice harboring the αC610S mutation would display cardiovascular pathologies.

Herein, we demonstrate that the KIC610S mice lacking GC1 transnitrosation activity and treated with Ang II display a higher increase in mean arterial pressure (MAP), compared to WT mice treated with Ang II.The KI mice also suffered from more severe cardiac hypertrophy, fibrosis, higher oxidation and electrical dysfunction in response to Ang II treatment, compared to WT. Surprisingly, the NO-stimulated GC1 activity did not decrease in the KI mice compared to the WT (± AngII). Further pressure myography and intravital microscopy studies in small resistance mesenteric arteries (MA) indicated that the KI mice has a significant decreased vasorelaxation in response to Acetylcholine (ACh) but vasorelaxation to NO remains essentially unchanged, even in the absence of oxidative stress. Overall these results suggest that GC1 transnitrosation activity modulates endothelium-dependent relaxation in small resistance vessels, hence regulates blood pressure.

## MATERIAL AND METHODS

### Generation of knock-in C610S mice

C610S knock-in mice were generated by the Genome Editing Shared Resource at Rutgers using CRISPR/Cas9 technology. The gRNA sequence to replace cysteine 610 with a serine was CTGCAGTGTGCCTCGGAAAATC →TAGCAGTGTGCCTCGGAAAATC and also included the restriction site BfaI (5’-C⋁TAG-3’) to facilitate screening of the Cys to Ser replacement, in addition to sequencing. 10 knock-in founders heterozygotes for C610S were successfully generated, 5 were bred (3 males, 2 females) with wild-type (WT) C57Bl/6J mice to expand the line. WT C57Bl/6 were purchased from Jackson laboratories and Charles River. The heterozygous C610S/+ first generation were backcrossed with WT C57Bl/6 and maintained on the C57Bl/6 genetic background. The homozygous WT and αGC1 C610S/C610S were generated by C610S/+ heterozygous breeding and whenever possible, littermates were used in the various experiments. 4 to 8-month-old male and female mice were used in the study. Mice were maintained in the animal facility at 22–24°C with 12-h light/dark cycles and 60%–70% of humidity. The animals were fed ad libitum under a normal chow diet and sterilized water. All animal experiments were conducted in accordance with policies of the NIH Guide for the Care and Use of Laboratory Animals and approved by the Institutional Animal Care and Use Committee (IACUC), Rutgers, Newark, NJ.

### HD-X11 telemetry device implantation and Angiotensin II (Ang II) infusion

Blood pressure (BP) and electrocardiogram (ECG) recordings were made in awake, freely moving mice using an implanted radio telemetry device (PhysioTel HD-X11 transmitter, Data Sciences International). Briefly, mice were anesthetized with 4-5% isoflurane inhalation, followed by single dose of local anesthetic Bupivacaine (2-2.5 mg/kg, SC). During surgery, anesthesia was maintained with 1-3% isoflurane during. The pressure catheter was inserted through the left carotid artery into the aortic arc, while the two leads were fixed subcutaneously in standard positions (positive lead was placed in left rib region and negative lead was placed to the right pectoral muscle).The transmitter body was secured into a subcutaneous pocket in the left flank. Buprenorphine (0.05 mg/kg, SC) was administered as an analgesic immediately after telemetry device implantation surgery and thereafter mice were allowed to recover in cages on heating Pad.

#### Angiotensin II (Ang II) infusion

At 5 to 7 days following telemetry device implantation, recordings were conducted as internal, untreated controls. At day 7, mice were treated with saline or AngII (0.8 mg/kg per day) for 14 days via a miniosmotic pump (ALZET^®^, cat # 0000296) implanted subcutaneously, as indicated in the scheme below.

#### Telemetry recordings

Mice were divided into 4 groups (WT ± Ang II and KI ± Ang II) and the data were recorded using a LabChart analysis software with a PhysioTel/HD Hardware MX2 configuration. Various pressures (systolic, diastolic and mean), heart rate (HR) and multiple ECG parameters, among them RR interval, PR interval, QT interval, QTc interval, QRS and P wave duration and amplitude were recorded and analyzed. The measurements were conducted in unrestrained mice, daily for 1 hour in the morning for 21 days. At the end of the experiment (day 28), mice were anesthetized and various organs including heart, lungs and mesenteric artery were removed, washed in PBS, and processed for biochemical or histological protocols.

#### Heart rate variability (HRV) analysis

HRV was performed on ECG tracings by using Labchart software. Arrhythmias, ectopic beats and artifacts were discarded from ECG tracing before HRV analysis by excluding R–R values not contained between mean R–R interval ± 2 S.D. (95.5% confidence intervals)(19).

#### Pressure Myography

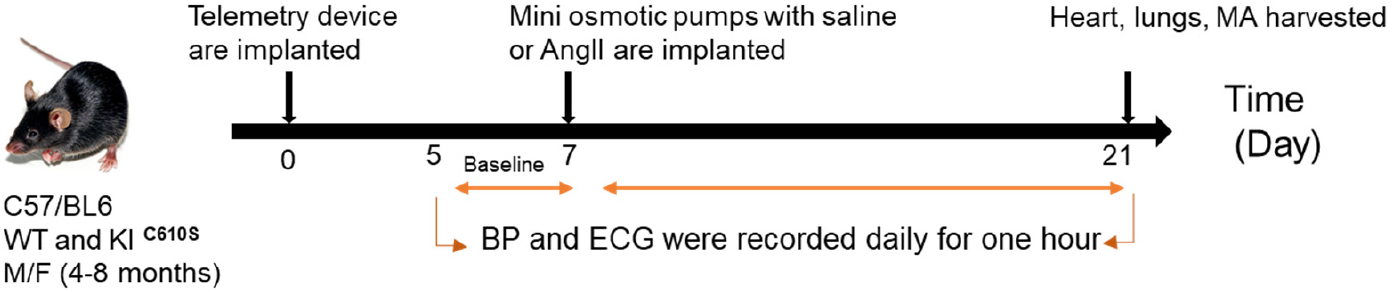
Schematic of the treatment schedule. C57/BL6 WT and KI ^C610S^ (male and female; 4-8 months old) were divided into 4 groups. WT ± Ang II and KI ^C610S^ ± Ang II. Telemetry device was implanted in the mice on day 0 and BP/ECG were recorded daily for one hour that served as baseline. Mini osmotic pumps with saline or Angll (0.8 mg/kg per day) were implanted on day 7 followed by daily one hour BP/ECG recordings until day 21. At day 21, mice were sacrificed, heart lungs and mesenteric arteries (MA) were harvested for further assays.

Mice were sacrificed under anesthesia and the mesenteric tissue was rapidly harvested and placed in a cold HEPES-PSS buffer containing (mmol/L): NaCl 118, KCl 4.7, CaCl_2_ 1.6, KH_2_PO_4_ 1.2, MgSO_4_ 1.2, NaHCO_3_ 25, glucose 5.5, HEPES 10. The final pH was adjusted to 7.4. For each experiment, a second or third order of mesenteric arteries (without any branches) was isolated and cannulated in a pressure myography chamber (Danish MyoTechnology A/S, model: 114P) using 60-80 or 100-120 µM cannulas. Each vessel was perfused in oxygenated prewarmed HEPES-PSS buffer (37°C, same composition that was used for vessel dissection). The pressure was increased gradually from 20 to 60 mmHg (with intervals of 10 mmHg). Bending of the vessel (due to pressure) was adjusted using the micro-positioners of the myography chamber; the vessel was equilibrated for 30 min at 37°C with a pressure of 60 mmHg (20). To evaluate the viability of vessel, 60 mM KCL solution was added to the chamber and vessels with less than 50% constriction were excluded from analysis. After KCl treatment, the chamber was washed 3 times with prewarmed HEPES buffer. The perfused vessel was live-tracked at 10 X magnification using a Zeiss AxioVert.A1 inverted microscope. Data including changes in outer and inner diameters were acquired and analyzed using the Myoview v. 5.1 software.

The vessels were pre-constricted with phenylephrine (PE at 10^−5^ M). Changes in vessel outer and inner diameters were then assessed in response to a cumulative dose of acetylcholine (ACh; 10^−8^-10^−4^ M) or Diethylamine NONOate diethylammonium (DEA/NO; 10^−8^-10^−4^ M). Percentage dilation was measured using the formula % 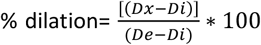, where D is the measured lumen diameter and x, i, and e indicate the measured arterial diameter at each dose of agonist (x), initial diameter following PE constriction (i), and diameter at 60 mmHg pressure after equilibrium (e) (20). n=3-7 mice per condition and 1-2 vessels were analyzed for each mouse.

### Vascular vasomotion response *in vivo* using intravital microscopy

To assess in vivo relaxation, we employed an intravital microscopy protocol as previously described (13, 21). Male and female mice, aged 4-6 months, were anesthetized with isoflurane (1.5-2%) for the procedure. The mesenteric bed was isolated for vessel visualization and continuously superfused with a saline solution at 37°C at a rate of 1 mL per minute. For vessel visualization and diameter detection, we inject Dextran 70-KDa-FITC conjugate retro-orbitally, as shown in **Supplemental Figure 1**. Vasodilators agents (10 µM ACh, 10 µM DEANO donor) were topically administered two minutes after 10 µM PE administration, during which time the peak and relative maintenance of arteriolar contraction were detected. The peak relaxation response was measured two minutes after administering ACh/NO, using the same formula as described in the pressure myography section. Diameter changes were quantified with Metamorph software (Molecular Device, USA). Relaxation was monitored using a time-lapse sequence, capturing images at one-second intervals. For each experimental condition, 2 to 3 vessels per animal were analyzed.

**Figure 1.**
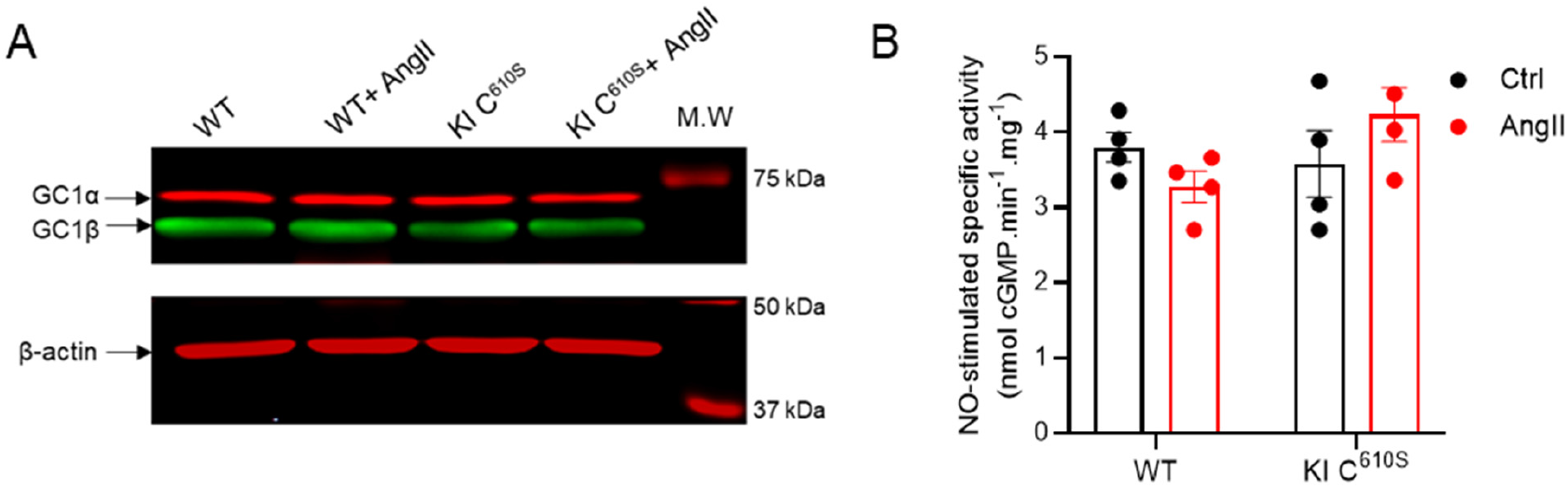
Assays of expression and NO-stimulated activity of WT and KI C610S mice ± Ang II. (**A**). Representative Western blot showing similar expression, albeit slightly less for the KI (n>3). Fifty μg of each lysate was electrophoresed on 12 % SDS gel then blotted and probed with antibodies against GC1α and GC1β subunits. β actin was used as loading control. (**B**) 10-50 μg lysates of lungs from the 4 groups of mice were assayed for basal (not shown) and NO-stimulated activity (the NO-donor DEA-NO was used at 100 μM). Lungs were used because they are one of the richest sources of GC1. n=4 mice. Measurements of GC1 activity were done in duplicate.

### Measurement of reactive oxygen species

We measured reactive oxygen species (ROS) levels in mice myocardial tissue with dihydroethidium (DHE), a redox-sensitive fluorescent probe (Cayman chemical company, cat # 104821-25-2). Briefly, frozen heart sections embedded with OCT (Tissue-Tek^®^, cat# 4583) were cut into 8-μm section, washed three times with PBS and incubated with DHE (2.5 µM) for 30 min at 37 °C in a dark humidified box. Following three washes with PBS, sections were incubated for 30 min at 37 °C with wheat germ agglutinin (WGA5µg/ml) to stain cardiomyocytes’ plasma membrane. Following three additional washes with PBS, sections were mounted in a medium-containing DAPI for nuclei staining (Sigma Aldrich, DUO82040). After drying in the dark, fluorescence images were captured using 200 Axiovert Zeiss fluorescence microscope, 20X objective and quantified with Image J software. To calculate DHE fluorescence in ImageJ, DAPI-stained nuclei were identified and corresponding fluorescence intensities of DAPI and DHE were measured. The fluorescence intensity of DHE was normalized to DAPI, and data are presented as the ratio of DHE/DAPI.

### Measurement of cardiac fibrosis

Heart tissues were fixed with 10% buffered formalin (Fisher chemical SF93-4), embedded in paraffin, and sectioned at 5μm thickness. Interstitial and perivascular fibrosis was evaluated by picric acid Sirius red staining. The sections were incubated with a 0.1% Sirius Red solution dissolved in aqueous saturated picric acid for 1 h, washed in acidified water (0.5% glacial acetic acid), dehydrated, and mounted with SignalStain medium (Cell Signaling, 14177S). Collagen is stained red, while non-collagen tissues are stained yellow. The positively stained (red) fibrotic area was measured and expressed as a percentage of total area. For each section, 6-10 fields at 10X magnification were used for the analysis by ImageJ. n=5-10 mice per condition.

### Soluble guanylyl cyclase activity measurements

Fresh or snap-frozen lung tissues (20-30 mg) were homogenized in a lysis buffer containing 50mM Tris-HCL pH 8.0, 150 mM NaCL, 1 mM EDTA and protease inhibitor cocktail + PMSF (Sigma, # P7626), using a Turax (VDI 12, VWR) on ice for 10-20 seconds. Homogenates were first spun at 4°C at 500g for 1 min to remove the biggest debris. The supernatant was then centrifuged at 14 000 g for 10 min at 4° C. The cytosolic fraction (supernatant) was used for Western blot and GC1 activity after measurement of protein concentrations using Bradford reagent. NO-stimulated GC1 activity was measured for 5 min. at 30° C in a 50mM HEPES pH 8.0 buffer containing 1mM GTP, 5 mM MgCl2, GTP^32^ and stimulated with 100 μM DEA-NO, as previously described (22). Each measurement was done in duplicate, n=4 mice for each group (WT, WT+ AngII, KI C^610S^, KI C^610S^ + AngII). Data are expressed in nmol cGMP.min^-1^.mg^-1^.

### Biotin switch assays of mouse tissues

Lung or heart tissues (20-30 mg) were homogenized in lysis buffer containing 50 mM Tris-HCL pH 7.5, 150 mM NaCl, 1 mM EDTA, 0.1 mM neocuproine, 0.2 mM PSMF and 40 mM N-ethylmaleimide (NEM). 2 mg of the homogenates were diluted to 0.5mg/ml in blocking buffer containing 1% Triton X-100 and 2% SDS in 50 mM Tris-HCL, pH 7.5, 150 mM NaCl, 1 mM EDTA, 0.1 mM neocuproine, 0.2 mM PSMF and 40 mM N-ethylmaleimide (NEM). Samples were incubated at 50°C for 30 min in the dark. Excess NEM was removed using cold acetone precipitation. The protein pellets were generated by centrifugation at 4000 rpm for 20 min at 4°C and washed once with cold acetone. The pellets were resuspended in 300 μl of a buffer containing 25 mM HEPES, pH 7.7, 1 mM EDTA, and 1% SDS, supplemented with 0.1 mM biotin-HPDP, 10 mM ascorbate and 1μM CuCl, at room temperature for 1 h in the dark. Negative controls were the blocked samples without ascorbate. Excess biotin-HPDP was removed and biotinylated proteins were precipitated using cold acetone at -20°C for 1 h followed by centrifugation at 5000 g for 10 min at 4°C. The pellets were dissolved in HENS buffer. For detection of biotinylated proteins, resuspended samples were mixed with 4X Laemmli buffer without β-Mercaptoethanol.

For avidin enrichment, the biotinylated samples were diluted in 700 μl PBS and mixed with 100 μl streptavidin-agarose beads. The mixture was incubated for 1 h at room temperature with regular agitation. The beads were pelleted by centrifuging at 5000 g for 10 min and washed 3 times with 1 ml of PBS. The washed beads were mixed with 100 μl of 1X Laemmli loading buffer with 10% β-Mercaptoethanol and heated at 85°C for 5 min. The proteins released from the beads were then subjected to Western blotting.

### Statistical analysis

Errors bars represent the mean ± SEM. Statistical analyses were performed using GraphPad Prism software (9.5.0). Two-way analysis of variance (ANOVA) followed by post-hoc Tukey’s multiple comparisons test were used when necessary. P values < 0.05 were considered significant for all statistical tests. Data collection and analyses were done blinded to the samples, whenever feasible.

## RESULTS

To determine the physiological significance of GC1-dependent transnitrosation activity, we created knock-in mice lacking this activity by replacing αC610, the main SNO donor cysteine of GC1, with a serine (αC^610S^). As previously, we used infusion of Angiotensin II to increase S-nitrosation and compromise vascular reactivity and cardiac homeostasis by augmenting oxidative stress (13, 14). Along the study, the 4 groups of mice were WT and KI αC^610S^ treated with AngII or vehicle. We first assayed whether the expression of GC1 and its activity were changed by the point mutation GC1-αC^610S^ compared to WT mice and in response to Ang II-induced oxidative stress.

### 1. The expression and NO-stimulated activity of GC1 are similar in WT and KI αC^610S^ mice ± AngII treatment

Lysates of lungs of the 4 groups of mice were prepared and the expression of GC1 α and β subunits assessed (Figure 1A). As previously observed (13, 14), the Ang II treatment did not affect GC1 expression, nor was the KI mutation, indicating that any difference in phenotype between WT and KI could not be attributed to a difference in expression. Likewise, the NO-stimulated activity of GC1 in lungs of WT and KI C^610S^ mice treated with vehicle or Ang II was not significantly different between the 4 groups (Figure 1B). As previously observed, Ang II treatment induces a decrease in NO-stimulated activity in the WT mice; however, it did not reach significance. The untreated KI C^610S^ mice has a NO-stimulated activity similar to untreated WT mice. In contrast to WT mice treated with AngII, the KI C^610S^ did not show any decrease in NO-stimulated activity when treated with AngII. Overall, while there might be a slight impairment of GC1 activity by oxidation in the WT, the expression and activities of GC1 in the 4 groups of mice are not altered by the point mutation C610S in the KI mice.

### 2. Ang II-induced global and specific S-nitrosation is drastically decreased in KI αC^s610S^ mice compared to WT mice

We next analyzed and compared to the WT mice, the S-nitrosation status of the KI C^610S^ mice under vehicle or Ang II treatment (Figure 2). For this, we conducted a biotin switch assay showing that biotinylation, hence nitrosation, was not detectable in the lung tissues of untreated WT and KI mice. In contrast, treatment with AngII led to high levels of S-nitrosation in WT mice but this increase was considerably blunted in KI mice (Figure 2A). Following avidin enrichment, specific S-nitrosation of GC1 was detected in WT and in KI mice (though with a slight decrease) with or without Ang II infusion. In the WT mice, S-nitrosation of RhoA and Trx1 were detected and increased with AngII. In sharp contrast, S-nitrosation was very faint in the KI mice whether they were treated or not with AngII. These results indicate that S-nitrosation of tissues in response to oxidative stress is largely modulated by GC1 and that αCys 610 is the main driver of GC1’ transnitrosation activity. As we previously observed in cells (6, 23) and now confirmed *in vivo*, GC1 is a major source of global S-nitrosation, S-nitrosated Trx1 and is also responsible for RhoA S-nitrosation probably via a transnitrosation cascade initiated by GC1.

**Figure 2.**
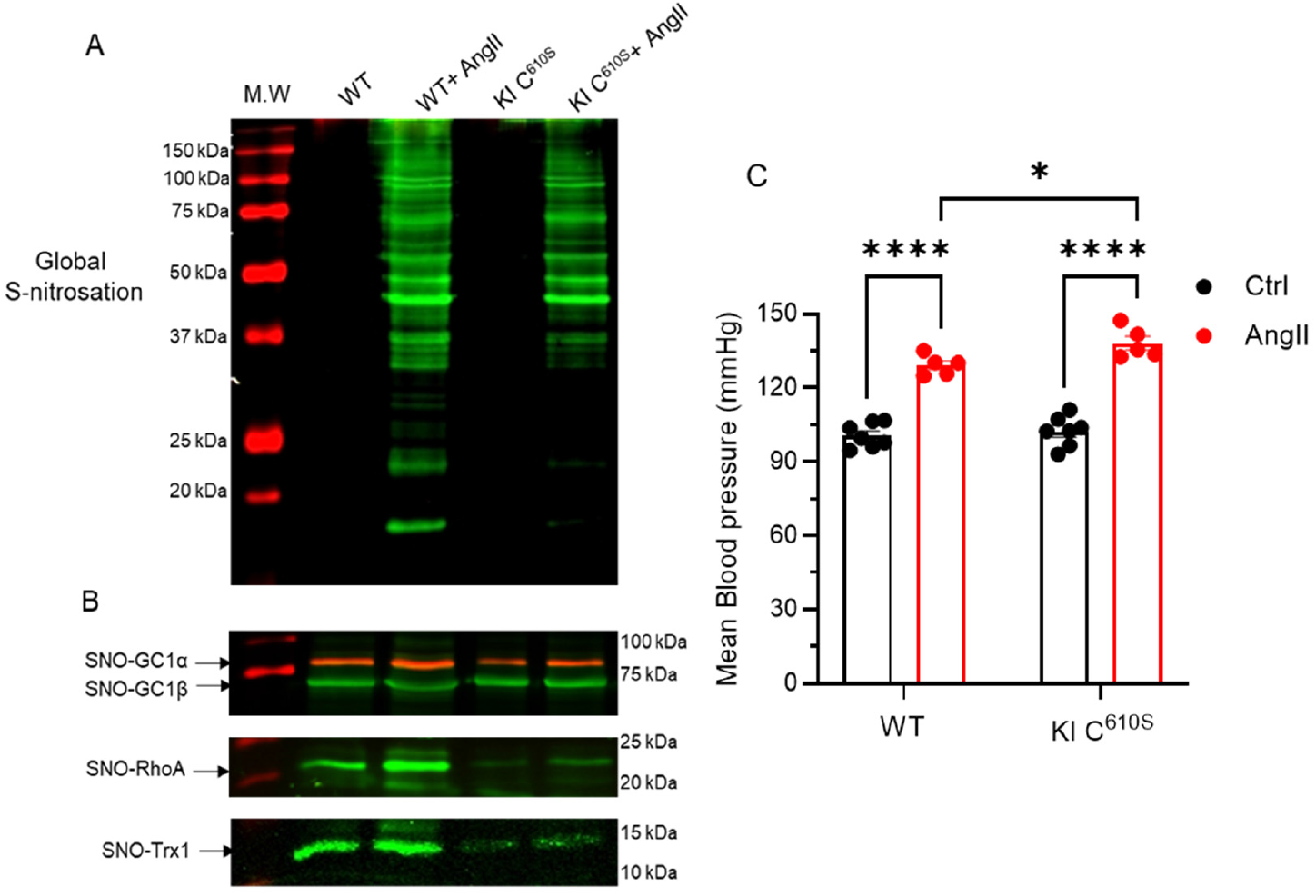
Decreased S-nitrosation in KI mice is associated with increased Ang II-induced blood pressure compared to WT. **Left panels**: S-nitrosation levels in WT and KI C610S mice ± Ang II. (**A**). Representative Western blot of biotinylated samples from the 4 groups electrophoresed on a non-reducing 12 % SDS-PAGE and probed with anti-biotin (*n*=3). (**B**) Following avidin purification, the biotinylated samples were separated by electrophoresis of a 12 % SDS-PAGE reducing gel, blotted and probed with antibodies against GC1 α and β, RhoA and Trx1. *n*=3. The uncut Ponceau red stained blots showing similar protein amount and the “no ascorbate” control are shown in **Supplemental Figure 2** for both Fig 2A and 2B. M.W.: Molecular Weight markers. **Right panel (C)**: Mean blood pressure (MBP) in WT and KI C610S mice ± Ang II. Telemetry recording of BP analyzed with LabChart software. n=5-7 for each group, expressed in mmHg ± SEM. Statistical analysis was performed with a two-way ANOVA-Tukey’s multiple comparisons test with *, p< 0.05, and ****, p< 0.001

### 3. AngII-dependent hypertension in WT mice is further increased in KI αC^610S^ mice

Next we aimed to test whether this deficit in S-nitrosation, in particular under AngII-induced oxidative stress, had any impact on the cardiovascular system. For this, we first assayed the changes in blood pressure. Using a telemetry system, we measured different pressures (systolic, diastolic and mean) in unrestrained mice, daily for 1 hour during a 21-day period. The telemetry device was implanted with a recovery period of 7 days and then the osmotic pump with AngII or vehicle was implanted for 14 days. Figure 2C (right panel) shows the mean blood pressure for each group at 7th day after the device implantation (vehicle, Ctrl) and at 14 days of AngII. Prior to AngII infusion, the baseline MBP measured at day 7 was similar between WT and KI, i.e., 105.9 ± 2.1 and 105.3 ± 3.3 mm Hg for the WT and KI mice, respectively. After 2 weeks of AngII treatment, telemetry recordings indicated significant increase in BP in both KI and WT mice, as expected(24) (25). However, the AngII-dependent rise in blood pressure was significantly higher (29.4%) in the KI mice compared to WT mice (20.9%), with a MBP of 130.6 ± 1.9 vs. 141.2 ± 3.3 mm Hg in the WT and KI mice, respectively. The increased pressure values in the KI mice imply that the cardiac output or/and the total resistance (vasorelaxation) is more affected by oxidative stress and correlate with decreased S-nitrosation in KI mice. This suggests that GC1-dependent transnitrosation activity could protect from cardiovascular oxidative pathologies.

### 4. Cardiac health is more compromised in KI mice treated with AngII than in WT mice treated with AngII

Because of a direct correlation between sustained elevated BP and cardiac health, we next assessed cardiac hypertrophy, oxidation, fibrosis and electrical properties in each group of mice. Figure 3A shows that there was no apparent cardiac hypertrophy, measured as the ratio of heart weight (mg) over body weight (g), in untreated WT and KI mice. AngII treatment induces hypertrophy in both WT and KI mice and was significantly higher in KI mice compared to WT mice treated with AngII (12.1% vs. 6.9% increase in KI and WT mice, respectively). As myocardial superoxide generation is a marker of Ang II-induced cardiac remodeling (26), we measured the levels of oxidation in the heart of the 4 groups of mice (Figure 3B and **supplementary Figure 3**). While DHE staining is increased by AngII treatment in both WT and KI mice, there was a significantly higher oxidation in KI mice in comparison to WT mice (ROS production was 1.5-fold higher in KI mice than WT mice after AngII treatment). These results indicate that reduced S-nitrosation in KI mice is associated with increased ROS and that the heart of KI mice is less protected from oxidative stress. Cardiac fibrosis is usually observed in Ang II-induced hypertension (27, 28). Using Picrosirius red staining of heart cryosections, we confirmed that AngII significantly increases interstitial and perivascular collagen deposition in both WT and KI mice. Importantly, KI mice exhibited more collagen deposition, hence more severe cardiac fibrosis than WT mice after AngII treatment (Figure 3C and **supplementary Figure 4**). The enhanced collagen deposition in KI mice suggests a potential protective effect of GC1-transnitrosation activity under oxidative stress conditions. To determine whether the cardiac fibrosis and hypertrophy were associated with electrical dysfunction, we conducted telemetric ECG recording in the same 4 groups of mice. While AngII treatment led to a prolonged QTc interval in both WT and KI, the increase in the duration was only significant in the KI mice (26.2% vs a 13.8% increase in WT + AngII). Consequently, the increase in QTc interval of KI + AngII mice was also significantly higher than in WT + AngII mice (Figure 4D). These data suggest that ventricular repolarization is delayed in KI mice. These results are consistent with previous studies that showed prolonged action potential durations in cardiomyocytes from hypertrophic hearts. The number of sinoatrial node (SAN) pauses was also significantly increased in KI mice treated with AngII; it was also higher compared to WT treated with AngII (Figure 4E), indicating a supraventricular dysfunction. Taken together, these results indicate that the KI mice develop hypertrophic cardiomyopathy accompanied with typical electrical alterations following AngII treatment. We further investigated the cardiac rhythm changes by extracting heart rate variability parameters (HRV) from the ECG recordings. We observed that the HRV was significantly decreased in KI mice compared to WT mice when challenged with Ang II (**supplementary Figure 5**).

**Figure 3:**
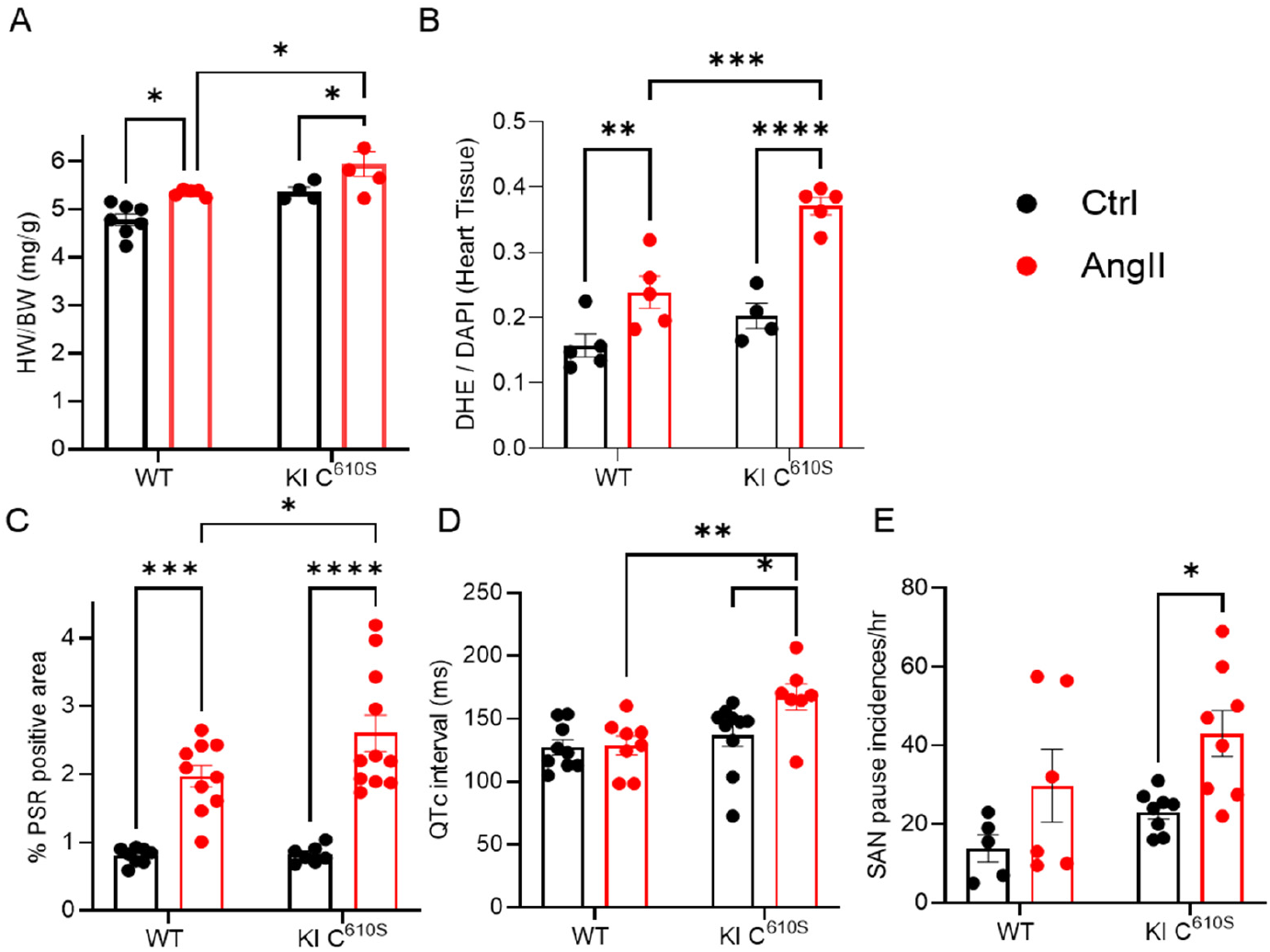
KI mice treated with AngII have more severe cardiac hypertrophy (A), higher cardiac oxidation (B), increased collagen deposition (C), longer QTc interval (D) and higher number of sinoatrial (SAN) pauses (E) compared to WT mice treated with AngII. Cardiac hypertrophy is expressed as HW (mg)/BW (g). Oxidation was assayed by DHE staining and fibrosis was assessed by Picrosirius red staining (PSR). Electrocardiogram (D and E) of the 4 groups of mice were recorded by telemetry and data were analyzed with labchart software. n=5-11 for each group. Values are expressed as S.E.M. Statistical analysis was performed with a two-way ANOVA-multiple comparisons test, with *, p< 0.05, **, p<0.01, and ****, p< 0.0001

**Figure 4:**
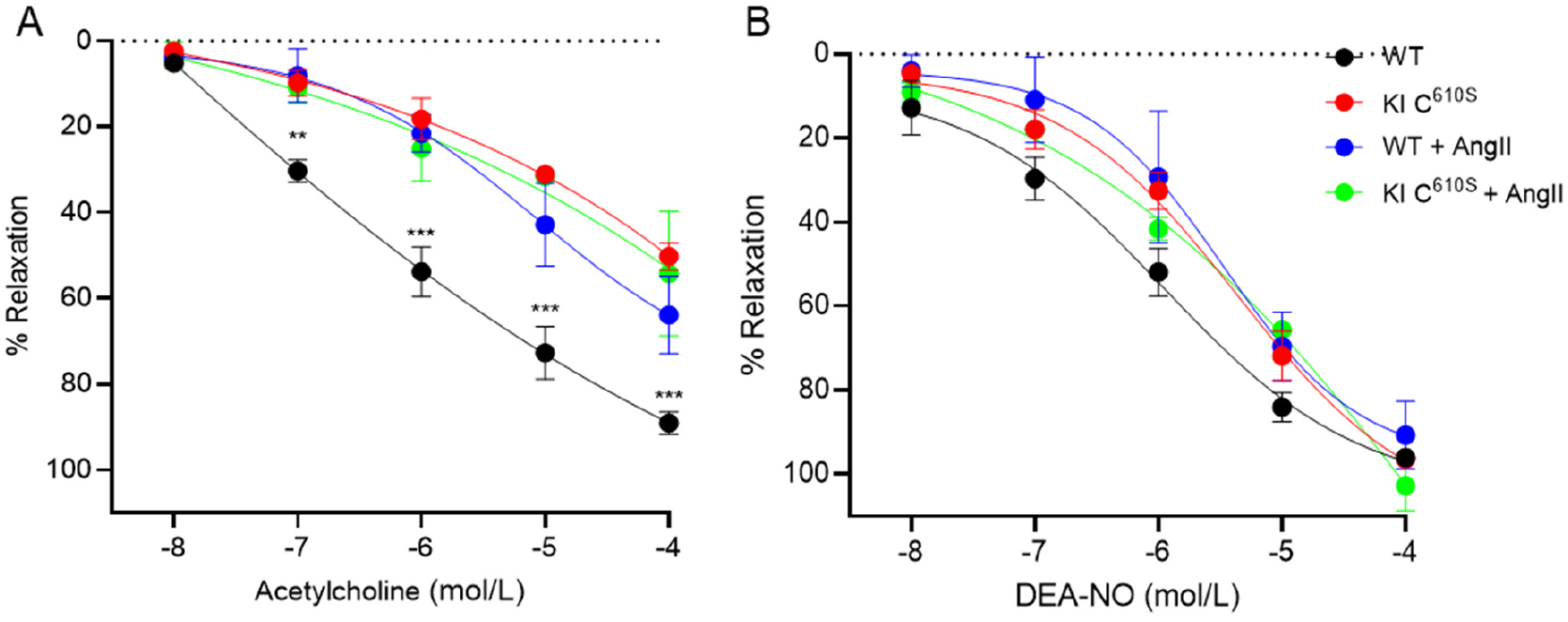
Cumulative ACh and DEA-NO dose-response curves of WT and KI C^610S^ mice untreated or treated with AngII. **(A)** Vasorelaxation to ACh was significantly impaired in the KI mice, as well as in the WT and KI treated with AngII, compared to untreated WT. There was no significant difference between KI, KI + AngII and WT + AngII. **(B)** The 4 groups showed a similar response to increasing DEA-NO concentrations. Vasorelaxation was assayed by changes in luminal diameter; data were recorded and analyzed with Myoview software. n=3 -6 for each group.. Statistical analysis was performed with a two-way ANOVA-Tukey’s multiple comparisons test, with **, p<0.01, and ***, p< 0.0001

**Figure 5.**
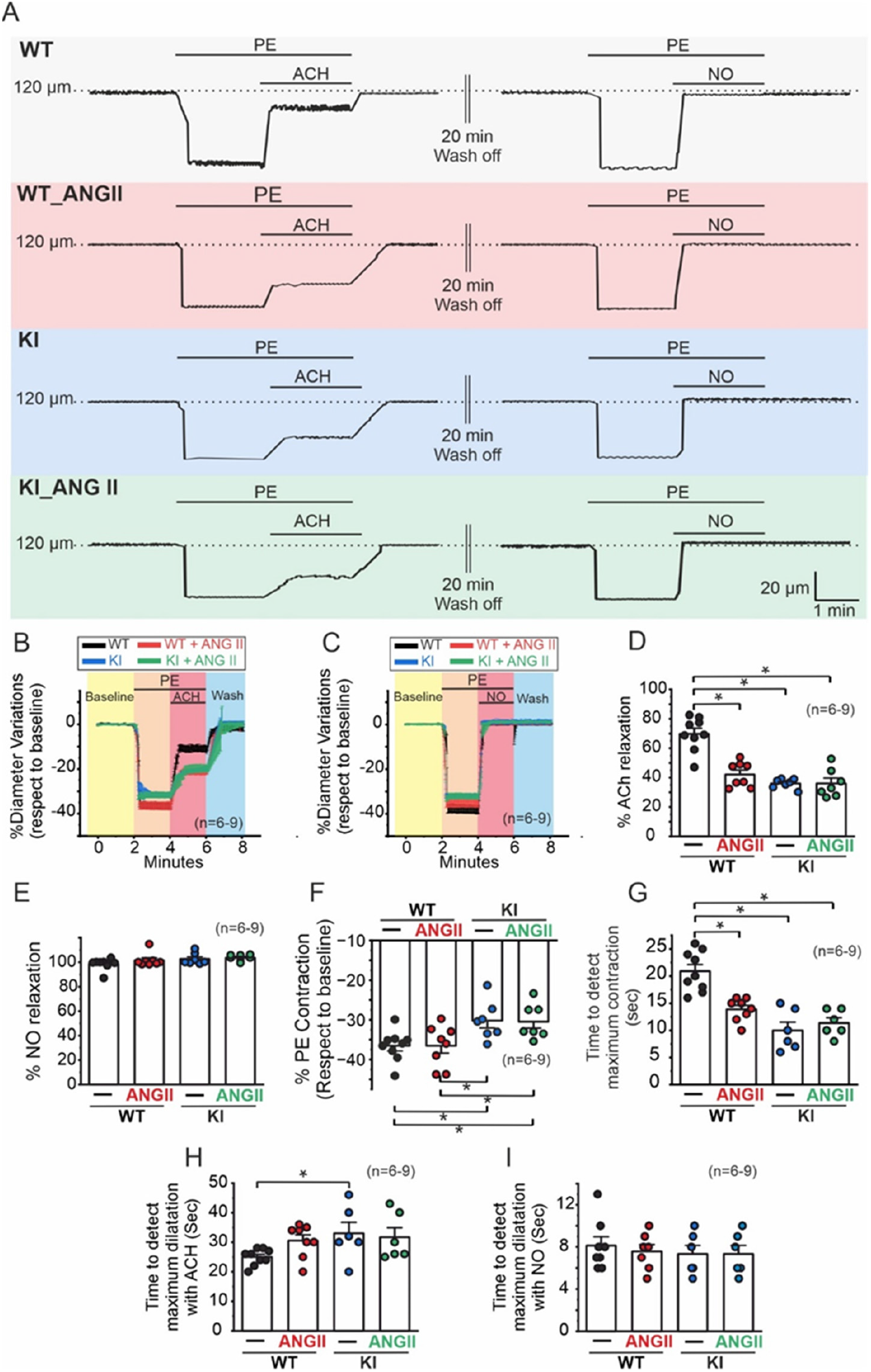
KI C610S mice exhibit diminished relaxation to ACh stimulation in PE-precontracted vessels *in vivo*. **A)** Representative traces showing vasomotion changes in the mesenteric bed of WT and KI mice under control conditions and AngII treatment. Vessels were stimulated with 10 µM PE, followed by 10 µM ACh or 10 µM DEANO to assess vasorelaxation. For each group of mice, average vascular response to ACh **(B)**, and to DEANO **(C)** were plotted (5-9 vessels, n>3 mice). **D)** Percentage of maximum relaxation response induced by ACh from data in B. **E)** Percentage of maximum relaxation response induced by DEA-NO, from data in C. From data in B, percentage of PE-induced contraction **(F)**, and time to achieve maximum contraction **(G)**. Time to achieve maximum dilation in PE-contracted vessels upon ACh **(H)** and upon DEA-NO **(I)**. Statistical analysis was performed with a two-way ANOVA-Tukey’s multiple comparisons test, with **, p<0.01

### 5. The endothelium-dependent relaxation in the resistance vasculature is significantly impaired in KI mice

To investigate the potential mechanism(s) of dysregulated BP in KI mice, we assessed the relaxation of the small resistance mesenteric arteries (MA) by pressure myography. The two main vasodilators of the resistance vessels are NO and endothelium hyperpolarization. NO induces relaxation of smooth muscle cells (SMC) through direct activation of GC1 (“muscle component” of relaxation), while hyperpolarization is usually considered the “endothelial component” of vasorelaxation. Acetylcholine (ACh) is the quintessential endothelial-dependent vasodilator, which triggers endothelial cell hyperpolarization in *small resistance arteries* leading to relaxation, while in *conduit (large) arteries* the ACh-induced NO release, in turn stimulating GC1 activity, is the predominant mechanism of vasorelaxation. We measured vascular resistance in isolated pressurized MA in the 4 groups of mice (WT, KI ± AngII) in response to ACh and the NO donor, DEA-NO. MA of 3-4^th^ order were pressurized at 60 mmHg, pre-contracted with 10 µM phenylephrine (PE) and increases in luminal diameter were assessed as a function of increasing (cumulative) concentrations of ACh and DEA-NO. We observed a significant decrease in ACh-dependent vasorelaxation in the KI mice compared to the WT. This decreased vascular response in the KI was similar in the WT and KI mice treated with AngII and significant compared to WT (Figure 4A). In sharp contrast, there was no significant changes in the vasorelaxation of the 4 groups of mice in response to DEA-NO, which directly stimulates GC1 activity in the SMC (Figure 4B). The lack of significant alteration in the response to NO in the resistance vasculature of the KI mice indicates that the NO-GC1-cGMP axis is not affected by the C610S replacement, in line with our observation that the levels of NO-stimulated production of cGMP remained unchanged in the KI mice. The impaired ACh-dependent vasorelaxation with maintenance of the NO-dependent response indicates that the C610S mutation does not affect the SMC component of vasorelaxation but leads to endothelium-dependent vascular dysfunction, probably at the origin of the increased BP in the KI mice. Importantly, the deficient response to ACh in the KI mice was observed in the absence of Ang II-induced oxidative stress suggesting that the mutation affects the physiological vasorelaxation signaling, even in the absence of oxidative stress.

#### 6-KI C610S Mice show reduced relaxation in PE-precontracted vessels in response to ACh stimulation *in vivo*

Figure 4 illustrates that the endothelial response is significantly diminished in KI mice with impaired S-nitrosation. To further evaluate these findings, we measured vasomotion *in vivo* in the mesenteric arteries, accounting for the influence of the surrounding microenvironment, such as adipocytes, sympathetic system and blood cells, among others. For this, we used intravital microscopy (**Supplemental Figure 1**). Under control conditions, arterioles ranging from 100 to 130 µm in outer diameter were precontracted with 10 µM PE. After two minutes of PE, these vessels were stimulated with 10 µM ACh or 10 µM DEA-NO. **Figure 5A** shows representative traces of the vasomotion response of arterioles from WT, WT + Ang II, KI, and KI + Ang II mice under baseline conditions and following PE, ACh, and NO-donor treatments.

Data analyses indicate that vasorelaxation peak in response to ACh was significantly impaired in KI mice as well as in Ang II-treated WT and KI mice compared to untreated WT (**Figure 5B and 5D**). In contrast, the response to NO remained unchanged in WT and KI mice untreated or treated with Ang II, aligning with pressure myography results that showed no difference in NO-smooth muscle relaxation (**Figure 5C and 5E**). Additionally, while the extent of contraction induced by PE was similar between WT mice with and without Ang II treatment, it was significantly reduced in KI mice under both conditions (control and Ang II, **Figure 5F**). Moreover, the time to reach peak contraction was significantly shorter (10-15 seconds) in KI mice and Ang II-treated WT mice, compared to WT mice for which the peak contraction typically occurs around 20-25 seconds. The shorter time to reach peak contraction in these models suggest faster smooth muscle contraction compared to WT mice (**Figure 5G**). Following ACh stimulation, the time required to achieve maximum relaxation was significantly prolonged in KI mouse models compared to WT mice. In contrast, this delay in relaxation was not observed with NO donor treatment (**Figures 5H and 5I**).

Collectively, the pressure myography and *in vivo* findings suggest that the endothelium-mediated vasorelaxation is specifically compromised by the C610S point mutation in the α-subunit of GC1. This mutation is linked to a reduction in GC1 transnitrosation activity while leaving the NO-GC1-cGMP signaling pathway intact. The fact that the defect in ACh-dependent relaxation takes place in the absence of oxidative conditions suggests that the mutation affects a physiological contractile signal.

## DISCUSSION

In this study, we established that the ability of the soluble guanylyl cyclase (GC1) to transfer S-nitrosothiol (SNO) groups, i.e., promotes S-nitrosation in other proteins, is involved in the vasorelaxation of small resistance vessels. This is an important finding as it indicates that the transnitrosation activity of GC1 contributes to regulation of blood pressure, which is largely determined by the peripheral vascular resistance. In the large conduit arteries, NO induces vasorelaxation by increasing cGMP production via stimulation of GC1 in the smooth muscle (SM) cells. On the other hand, vasorelaxation of small resistance vessels is initiated by endothelium hyperpolarization, which in turn relaxes SM cells. While the identity of endothelium–derived hyperpolarization factors and the exact mechanism(s) are still highly debated, S-nitrosation appears to be one of the contributing factors of endothelium-dependent vasorelaxation(3, 29-31). Acetylcholine (ACh) the quintessential endothelium-dependent vasodilator also activates endothelial NO synthases leading to NO production and subsequent activation of GC1 in the SMC. For this reason it has been very difficult to distinguish the contribution of the endothelium vs. SM component of vasorelaxation regulated by the NO/SNO vasodilation pathways. By introducing in mice a single mutation of GC1 (C610S) that affects its transnitrosation activity but leaves intact its “canonical” NO-stimulated cGMP forming activity, we were able to uncouple two keys NO signaling pathways that are involved in the endothelial and SM components of vasorelaxation.

We have shown that the NO-stimulated activity of GC1 is inhibited by S-nitrosation under oxidative stress. Many reports demonstrated that excess S-nitrosation was associated with endothelial dysfunction(13, 32). When we treated the WT and KI C610S mice with Ang II, our intent was to determine whether the mutation, by potentially lowering the levels of S-nitrosation in the KI, will be protective from extensive cardiovascular pathologies. The opposite took place as shown by exacerbated cardiac pathologies and a higher rise in MAP in the KI under Ang II treatment. Remarkably, the NO-cGMP axis did not appear to be involved, because the mutation did not decrease NO-stimulated cGMP activity and because NO-dependent vasorelaxation of SMC still occurred in the isolated MA and *in vivo*. In contrast, the ACh-dependent vasorelaxation in the resistance vessels was strongly and significantly decreased and the alteration was maximal in the KI, even in the absence of AngII treatment. One potential explanation was that the mutation C610S, considering its overall effect on global S-nitrosation levels removes an attributed function of SNO, which is to protect the tissues from more deleterious thiol oxidations, in particular in the cardiac tissues (15). This could be the explanation for the compromised cardiac heath observed in the KI mice treated with AngII. Nonetheless, we observed the decreased ACh-dependent relaxation in the absence of oxidative/nitrosative stress suggesting that the mutation C610S impaired a cGMP-independent physiological function of GC1, probably unrelated to a mechanism of protection in oxidative pathologies. We did not observe an increased MAP in untreated KI mice and inferred that this could be due to compensatory mechanism(s) from the multiple regulators of BP.

In summary, we propose that GC1-dependent transnitrosation is involved in the endothelium– dependent vasorelaxation of resistance vessels and our next challenge is to identify and assess the involvement of SNO targets, known to participate in endothelium-dependent hyperpolarization mechanism. We also currently investigate the role of RhoA in these phenotypes because it is a GC1/Trx1 transnitrosation cascade target *in vivo*, and its S-nitrosation leads to inhibition of its contractile pathway(33). Likewise, we will assess whether C610 is the potential cys that was proposed to interact with additional NO to maximally stimulate GC1, a somewhat controversial mechanism (34) (addition of excess NO in the pressure myography and *in vivo* experiments might have masked this mechanism). Likewise, the impact on cardiac pathologies of the mutation C610S and the underlying mechanisms need to be further explored (5).

## ACKNOWLEDGMENTS

This work was supported by the National Institutes of Health, R01 GM 067640 and R01 GM112415 (A.B.); R01 HL157116 and R01HL133294 (L.H.X). American Heart Association, 19TPA34900003 (L.H.X.) Career Development Award 932684 (M.L.); AHA Research Supplement to Promote Diversity in Science 23DIVSUP1054931 (P.B.).

## REFERENCES

1. J. Erdmann et al., Dysfunctional nitric oxide signalling increases risk of myocardial infarction. Nature 10.1038/nature12722 (2013).

2. E. J. Tsai, D. A. Kass, Cyclic GMP signaling in cardiovascular pathophysiology and therapeutics. Pharmacol Ther 122, 216–238 (2009).

3. Y. Zhao, P. M. Vanhoutte, S. W. Leung, Vascular nitric oxide: Beyond eNOS. J Pharmacol Sci 129, 83–94 (2015).

4. P. Sandner et al., Soluble Guanylate Cyclase Stimulators and Activators. Handb Exp Pharmacol 264, 355–394 (2021).

5. C. T. Stomberski, D. T. Hess, J. S. Stamler, Protein S-Nitrosylation: Determinants of Specificity and Enzymatic Regulation of S-Nitrosothiol-Based Signaling. Antioxid Redox Signal 10.1089/ars.2017.7403 (2018).

6. C. Cui et al., Soluble guanylyl cyclase mediates noncanonical nitric oxide signaling by nitrosothiol transfer under oxidative stress. Redox biology 55, 102425 (2022).

7. R. C. Shah, S. Sanker, K. C. Wood, B. G. Durgin, A. C. Straub, Redox regulation of soluble guanylyl cyclase. Nitric Oxide 76, 97–104 (2018).

8. E. Schulz, T. Gori, T. Munzel, Oxidative stress and endothelial dysfunction in hypertension. Hypertens Res 34, 665–673 (2011).

9. E. S. Arner, A. Holmgren, Physiological functions of thioredoxin and thioredoxin reductase. Eur J Biochem 267, 6102–6109 (2000).

10. S. I. Hashemy, A. Holmgren, Regulation of the catalytic activity and structure of human thioredoxin 1 via oxidation and S-nitrosylation of cysteine residues. J Biol Chem 283, 21890–21898 (2008).

11. C. Wu et al., Thioredoxin 1-Mediated Post-Translational Modifications: Reduction, Transnitrosylation, Denitrosylation, and Related Proteomics Methodologies. Antioxid Redox Signal In press (2011).

12. C. Huang et al., Guanylyl cyclase sensitivity to nitric oxide is protected by a thiol oxidation-driven interaction with thioredoxin-1. J Biol Chem 292, 14362–14370 (2017).

13. P. A. Crassous et al., Soluble guanylyl cyclase is a target of angiotensin II-induced nitrosative stress in a hypertensive rat model. American journal of physiology. Heart and circulatory physiology 303, H597–604 (2012).

14. P. A. Crassous et al., Newly Identified NO-Sensor Guanylyl Cyclase/Connexin 43 Association Is Involved in Cardiac Electrical Function. Journal of the American Heart Association 6 (2017).

15. E. Murphy, M. Kohr, J. Sun, T. Nguyen, C. Steenbergen, S-nitrosylation: a radical way to protect the heart. J Mol Cell Cardiol 52, 568–577 (2012).

16. J. Sun, M. Morgan, R. F. Shen, C. Steenbergen, E. Murphy, Preconditioning results in S-nitrosylation of proteins involved in regulation of mitochondrial energetics and calcium transport. Circ Res 101, 1155–1163 (2007).

17. B. Lima, M. T. Forrester, D. T. Hess, J. S. Stamler, S-nitrosylation in cardiovascular signaling. Circ Res 106, 633–646 (2010).

18. I. H. Schulman, J. M. Hare, Regulation of cardiovascular cellular processes by S-nitrosylation. Biochim Biophys Acta 1820, 752–762 (2012).

19. J. Thireau, B. L. Zhang, D. Poisson, D. Babuty, Heart rate variability in mice: a theoretical and practical guide. Exp Physiol 93, 83–94 (2008).

20. M. Shahid, E. S. Buys, Assessing murine resistance artery function using pressure myography. Journal of visualized experiments : JoVE 10.3791/50328 (2013).

21. N. Sayed et al., Nitroglycerin-induced S-nitrosylation and desensitization of soluble guanylyl cyclase contribute to nitrate tolerance. Circ Res 103, 606–614 (2008).

22. A. C. Straub, A. Beuve, A primer for measuring cGMP signaling and cGMP-mediated vascular relaxation. Nitric Oxide 117, 40–45 (2021).

23. C. Cui et al., Inhibitory Peptide of Soluble Guanylyl Cyclase/Trx1 Interface Blunts the Dual Redox Signaling Functions of the Complex. Antioxidants (Basel) 12 (2023).

24. K. Okuno et al., Angiotensin II Type 1A Receptor Expressed in Smooth Muscle Cells is Required for Hypertensive Vascular Remodeling in Mice Infused With Angiotensin II. Hypertension 80, 668–677 (2023).

25. J. C. Romero, J. F. Reckelhoff, State-of-the-Art lecture. Role of angiotensin and oxidative stress in essential hypertension. Hypertension 34, 943–949 (1999).

26. E. Takimoto, D. A. Kass, Role of oxidative stress in cardiac hypertrophy and remodeling. Hypertension 49, 241–248 (2007).

27. K. Broekmans, J. Giesen, L. Menges, D. Koesling, M. Russwurm, Angiotensin II-Induced Cardiovascular Fibrosis Is Attenuated by NO-Sensitive Guanylyl Cyclase1. Cells 9 (2020).

28. J. M. Schnee, W. A. Hsueh, Angiotensin II, adhesion, and cardiac fibrosis. Cardiovasc Res 46, 264–268 (2000).

29. S. M. Mutchler, A. C. Straub, Compartmentalized nitric oxide signaling in the resistance vasculature. Nitric Oxide 49, 8–15 (2015).

30. Y. Iwakiri, S-nitrosylation of proteins: a new insight into endothelial cell function regulated by eNOS-derived NO. Nitric Oxide 25, 95–101 (2011).

31. S. Luo et al., Roles of Protein S-Nitrosylation in Endothelial Homeostasis and Dysfunction. Antioxid Redox Signal 40, 186–205 (2024).

32. H. Choi, K. J. Allahdadi, R. C. Tostes, R. C. Webb, Augmented S-nitrosylation contributes to impaired relaxation in angiotensin II hypertensive mouse aorta: role of thioredoxin reductase. J Hypertens 29, 2359–2368 (2011).

33. L. Lin et al., RhoA inactivation by S-nitrosylation regulates vascular smooth muscle contractive signaling. Nitric Oxide 74, 56–64 (2018).

34. N. B. Fernhoff, E. R. Derbyshire, M. A. Marletta, A nitric oxide/cysteine interaction mediates the activation of soluble guanylate cyclase. Proc Natl Acad Sci U S A 106, 21602–21607 (2009).

