## Supplementary information for "Soluble guanylyl cyclase, the NO-receptor, regulates endothelium-dependent vascular relaxation via its transnitrosation activity"

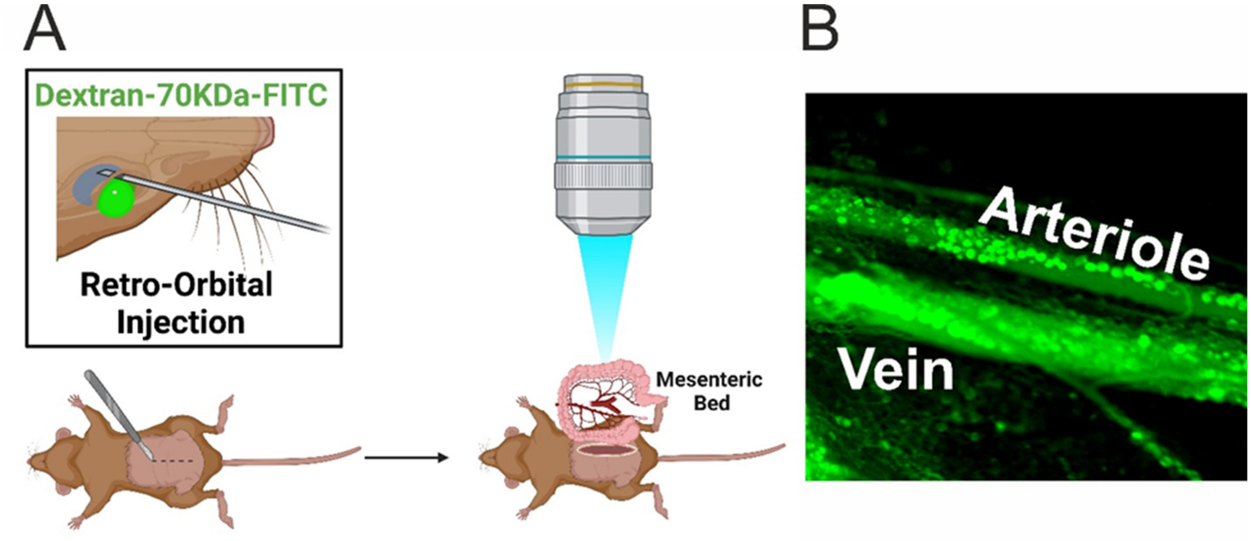


**Supplemental Figure 1. Intravital Microscopy Method for Assessing Vascular Vasomotion Changes. A)** Mesenteric vessels are visualized using intravital microscopy **in vivo** following retro-orbital injection of Dextran 70-KDa-FITC conjugate. The mesenteric bed is then exposed and perfused with saline solution at 37°C. Imaging is performed with a 5x long-distance objective (Leica N Plan 5x Objective) to capture the microvasculature. **B)** Representative images of the mesenteric microvascular bed are shown, highlighting both arterioles and veins. Changes in vasomotion in arterioles, induced by vasoactive agents, are monitored and analyzed using Metamorph software.

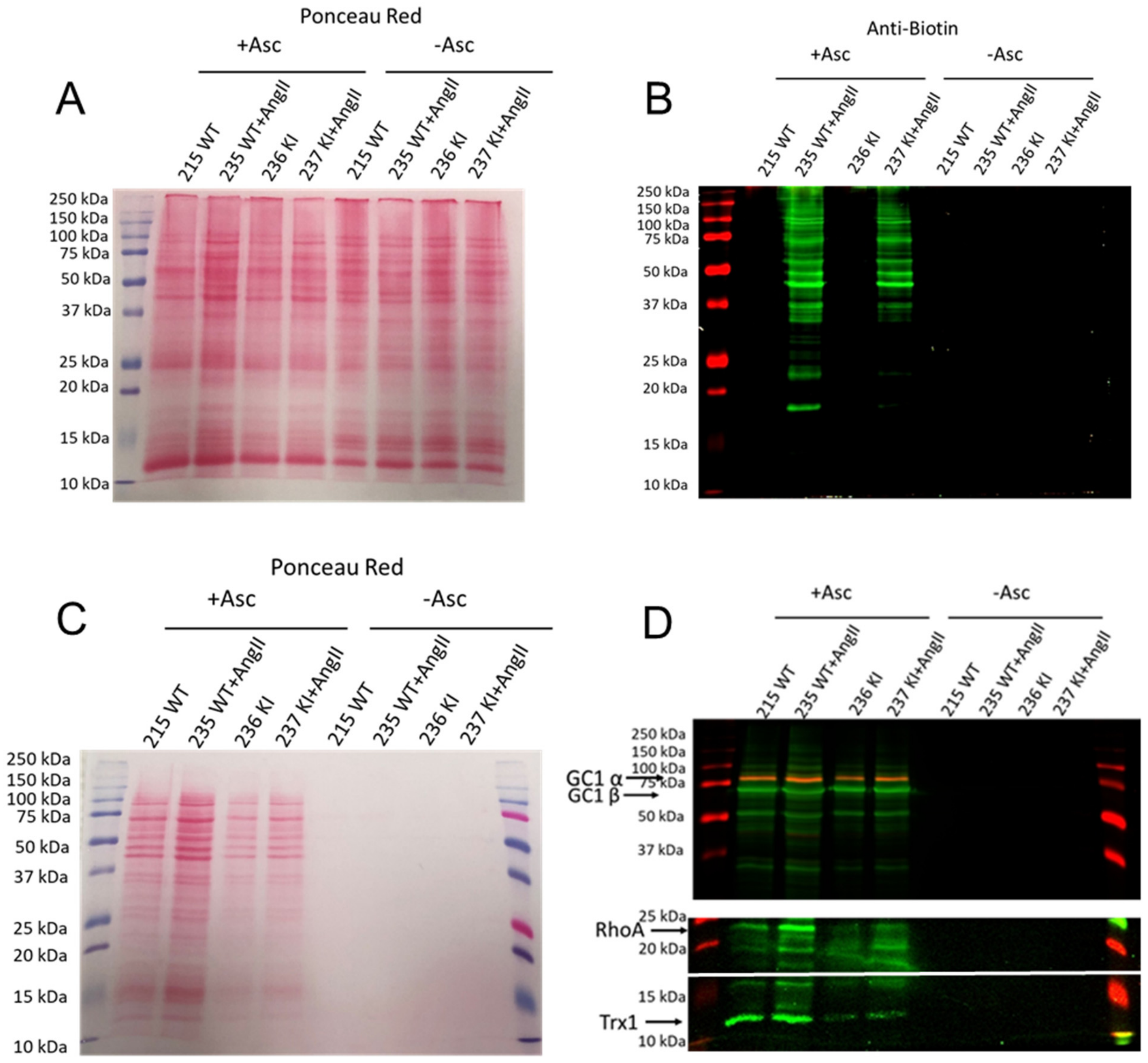


**Supplemental Figure 2. Ponceau red and Western blots from representative Fig. 2A and 2B.**
Ponceau red of membrane with biotinylated samples (A), probed with anti-biotin (B)
Ponceau red of membrane with avidin-enriched samples (C), probed with anti-GC1α, anti-GC1β (upper panel), anti-RhoA and anti-Trx1 (lower panel).

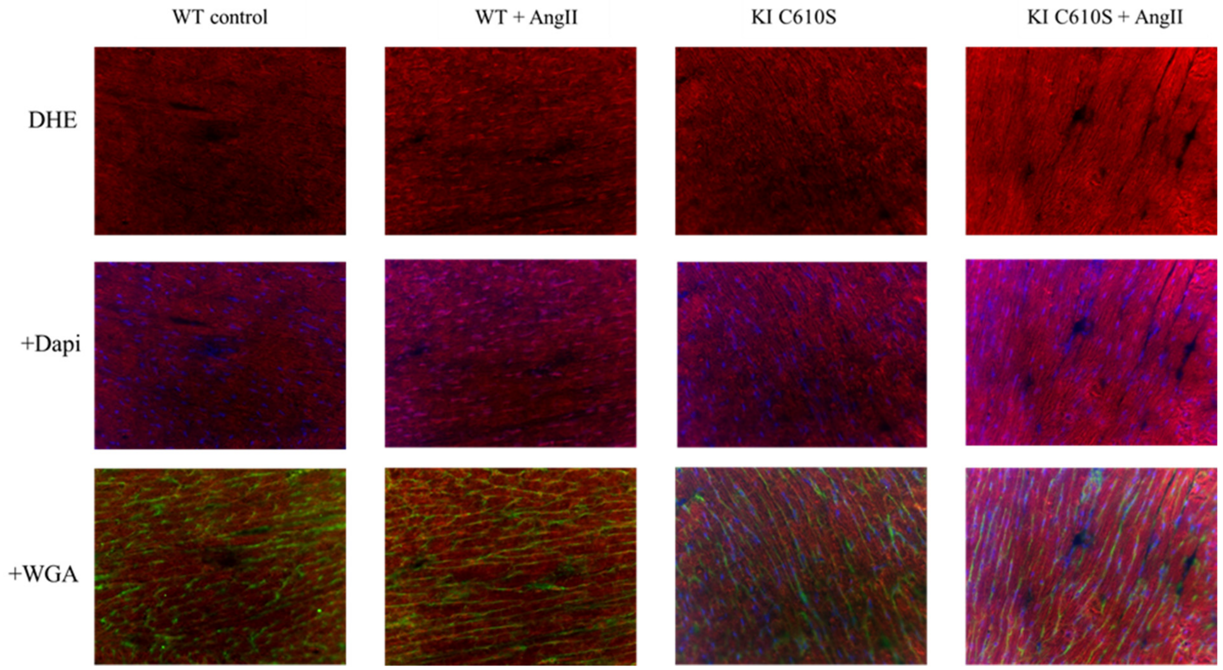


**Supplemental Figure 3**. Representative images showing *in situ* detection of ROS production with dihydroethidium (DHE; red fluorescence) in heart cryosections (8 μm) of WT and KI $^{C610S}$ mice with and without AngII treatment. Fluorescence images were captured using 200 Axiovert Zeiss microscope, 20x objective and quantified with Image J software. Dapi in the mounting medium was used to stain the nuclei in blue; WGA: Wheat Germ Agglutinin to stain membranes (green).

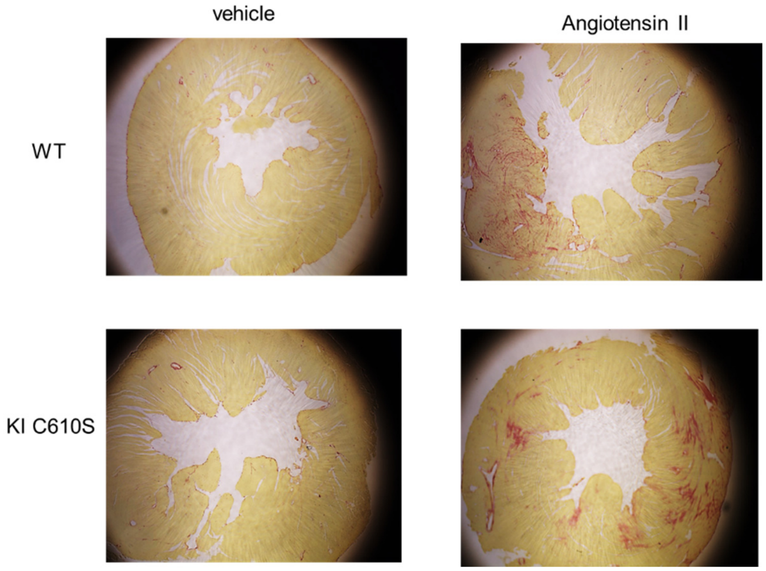


**Supplemental Figure 4**. Representative images : Representative images of staining with picric acid/red Sirius of WT and KI mice heart sections treated with vehicle of angiotensin II. Objective 2 X magnification. (collagen is stained in red).

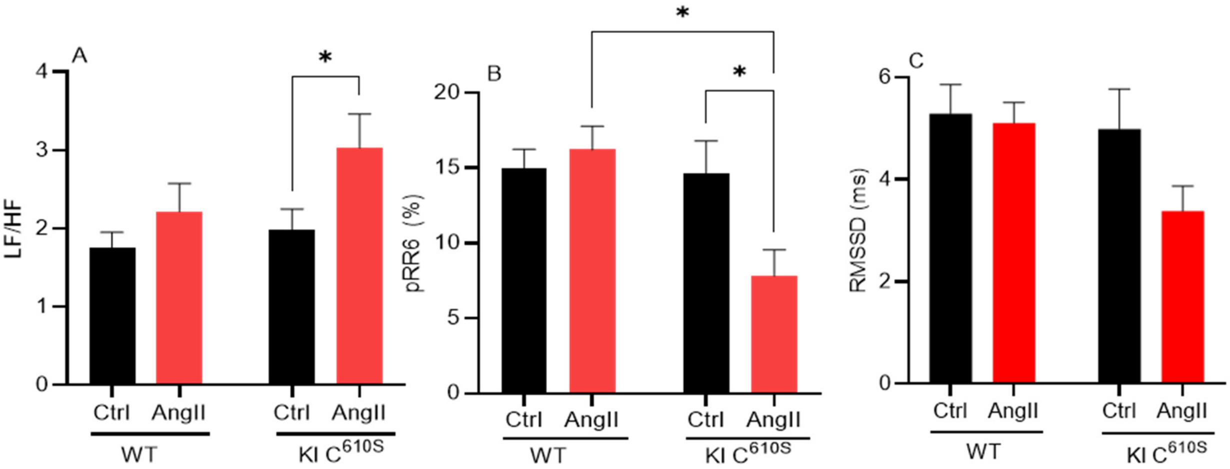


**Supplementary Fig.5**: Ang II-treated KI mice have decreased heart rate variability (HRV) as indicated by higher LF/HF ratio (A), lower pRR6 % (B), and decreased RMSSD (C) compared to controls and/or WT treated with AngII. The ECG of the 4 groups of mice analyzed with the module HRV of labchart. n=5. LH and HF: Low and High frequency power, respectively; pRR6: percentage of normal consecutive R-R intervals differing by > 6 ms. RMSSD: Root Mean Squared of Successive differences. Statistical analysis was performed with a two-way ANOVA-Tukey's multiple comparisons test; *, $p < 0.05$; **, $p < 0.01$, and ***, $p < 0.001$.
